# Contextual fear learning and memory differ between stress coping styles in zebrafish

**DOI:** 10.1101/535294

**Authors:** Matthew R Baker, Ryan Y Wong

**Affiliations:** Department of Biology, University of Nebraska at Omaha, Omaha, Nebraska, USA; Department of Psychology, University of Nebraska at Omaha, Omaha, Nebraska, USA

**Keywords:** Animal Personality, Stress Coping Style, Cognitive Biases, Learning and Memory, Alarm Substance, Zebrafish

## Abstract

Animals frequently overcome stressors and the ability to learn and recall these salient experiences is essential to an individual’s survival. As part of an animal’s stress coping style, behavioral and physiological responses to stressors are often consistent across contexts and time. However, we are only beginning to understand how cognitive traits can be biased by different coping styles. Here we investigate learning and memory differences in zebrafish (*Danio rerio*) displaying proactive and reactive stress coping styles. We assessed learning rate and memory duration using an associative fear conditioning paradigm that trained zebrafish to associate a context with exposure to a natural olfactory alarm cue. Our results show that both proactive and reactive zebrafish learn and remember this fearful association. However, we note significant interaction effects between stress coping style and cognition. Zebrafish with the reactive stress coping style acquired the fear memory at a significantly faster rate than proactive fish. While both stress coping styles showed equal memory recall one day post-training, reactive zebrafish showed significantly stronger recall of the conditioned context relative to proactive fish four days post-training. Through understanding how stress coping strategies promote biases in processing salient information, we gain insight into mechanisms that can constrain adaptive behavioral responses.

## Introduction

When animals successfully overcome stressors, cognitive processes facilitate the encoding and recalling of these salient experiences to modify or reinforce beneficial coping behaviors in the future. Within an individual, behavioral and physiological responses to stress often co-vary as part of a correlated suite of traits that are consistent across contexts and time (i.e. animal personality)(Baker et al., 2017; Koolhaas et al., 1999; Koolhaas et al., 2010; Øverli et al., 2007). Animals that are risk-prone or risk-averse differ in boldness, aggression, and stress physiology, and represent opposite ends of a response continuum observed across many taxa (e.g. bold-shy, proactive-reactive axis)(Koolhaas et al., 1999, 2010; Øverli et al., 2007; Sih et al., 2004). While variation in cognitive abilities can be due to a variety of factors (Dalesman, 2018; Lucon-Xiccato & Bisazza, 2017; Miller, 2017; Sorato et al., 2018), studies are beginning to demonstrate that learning and memory processes are also biased according to personality type (Brown et al., 2013; Dougherty & Guillette, 2018; Lucon-Xiccato & Bisazza, 2017; Miller, 2017; Sih & Del Giudice, 2012).

In line with other behavioral and physiological traits, studies suggest that proactive and reactive stress coping styles differ in information processing, decision making, and learning and memory capabilities (Carere & Locurto, 2011; Dougherty & Guillette, 2018; Griffin et al., 2015; Lucon-Xiccato & Bisazza, 2017; Øverli et al., 2007; Sih & Del Giudice, 2012). The more risk-prone proactive individuals tend to rely on past experiences and form more rigid routines (i.e. low behavioral flexibility). In contrast, the risk-averse reactive individuals are more sensitive to environmental cues for learned associations and display higher behavioral flexibility. Despite these observations, there are inconsistencies across studies investigating how learning and memory abilities vary with personality type in mammals, birds, and teleosts, often relating to the type of paradigm and stimulus valence. Some studies show that reactive individuals will learn faster (Budaev & Zhuikov, 1998; Exnerová et al., 2010; Miller et al., 2006), but others show support for proactive individuals learning faster (Amy et al., 2012; Bolhuis et al., 2004; DePasquale et al., 2014; Dugatkin & Alfieri, 2003; Mazza et al., 2018; Mesquita et al., 2015; Trompf & Brown, 2014). The same conflicting observations are documented with memory performance between the stress coping styles (Brown et al., 2013; Exnerová et al., 2010; Moreira et al., 2004). Examining to what extent encoding and recalling of salient information is influenced by stress coping style is important towards understanding factors that may facilitate the development of correlated suites of traits within an individual.

Exposure to highly stressful events such as predation are useful for investigating individual differences in learning and memory. Upon experiencing a threatening event, an individual can associate a specific cue of the threatening stimulus and the general environment in which it was experienced (e.g. context)(Maren et al., 2013). Many learning paradigms utilize predator odors or chemical alarm signals as an unconditioned stimulus (US) to study ecologically relevant cognitive behaviors (Takahashi et al., 2008). In teleosts a chemical alarm signal (alarm substance) is released from epidermal cells when they are mechanically damaged. This olfactory signal causes robust antipredatory behaviors even in the absence of a predator, and is used to assess stress-related behaviors in zebrafish (*Danio rerio*) and other teleosts (Gerlai, 2010; Speedie & Gerlai, 2008). Typical fear responses in teleost include bottom dwelling, swimming in a tighter shoal, erratic movements and freezing. While studies have utilized alarm substance for associative conditioning paradigms of specific cues on schools of fish, it has presented some challenges for measuring individual differences in learning and memory (Brown et al., 2013; Hall & Suboski, 1995; Ruhl et al., 2017). Further not much is known whether alarm substance can be used for contextual learning and recall of salient information. Utilizing alarm substance to study the relationship between learning, memory, and personality types will require behavioral assays that can be tested on individual fish, are rapidly and reliably acquired, and allow for isolated examination of both learning and memory recall phases.

Here we test for differences in how contextual associations are formed and maintained between two lines of zebrafish selectively bred to display proactive and reactive stress coping styles in an associative fear conditioning task. Using a novel contextual fear conditioning paradigm, we compared the rate fish learned to associate a formerly neutral context with a fearful antipredatory response induced by exposure to alarm substance. Additionally, we tested memory recall at two different time points following training to assess the duration of fear memory retention.

## Methods

### Subjects

Here we use zebrafish to study how cognitive abilities varies with stress coping style. Zebrafish are utilized in a variety of laboratory studies to understand the neural, genetic, and pharmacological mechanisms of learning and memory (Gerlai, 2016; Norton & Bally-Cuif, 2010; Oliveira, 2013). Both wild and laboratory strains of zebrafish display the proactive and reactive stress coping styles, which have distinct genetic architectures and neuroendocrine responses (Oswald et al., 2012; Oswald et al., 2013; Russ, 2018; Wong et al., 2015). Given their rich repertoire of learning and memory behaviors, low costs, high-throughput assays, genetic tractability, evolutionary significance, and homologous anatomy and physiology to their mammalian counterparts, zebrafish are a promising system to study how an animal’s stress coping style influences fear learning and memory abilities (Bshary & Brown, 2014; Gaikwad et al., 2011; Gerlai, 2010; Ijaz & Hoffman, 2016; Norton & Bally-Cuif, 2010; Oliveira, 2013).

We specifically used the high-stationary behavior (HSB) and low-stationary behavior (LSB) zebrafish strains (Wong et al., 2012). Starting from wild-caught zebrafish, the HSB and LSB strains were generated and are maintained by artificial selection for opposing amounts of stationary behavior to a novelty stressor (Wong et al., 2012). The HSB and LSB strains show contrasting behavior, physiology, morphology, and neuromolecular profiles consistent with the reactive and proactive coping styles, respectively (Kern et al., 2016; Russ, 2018; Wong & Godwin, 2015; Wong et al., 2015; Wong et al., 2014; Wong et al., 2013). Additionally, these divergent behavioral profiles between the strains are consistent across contexts and over time and are highly repeatable (Baker et al., 2018; Wong et al., 2012). We tested 32 individuals for each of the LSB and HSB strains. Fish that did not display any response to the US were removed from the study, resulting in a final sample size of 24 LSB (N = 12 males, 12 females) and 24 HSB (N = 12 males, 12 females) for the treatment group receiving alarm substance during training. An additional 8 LSB (N = 4 males, 4 females) and 8 HSB (N = 4 males, 4 females) were used as a control group being exposed to distilled (DI) water during training. LSB and HSB individuals were 16 months post-fertilization when testing began. During testing, fish were individually housed in 3-liter tanks on a recirculating water system (Pentair Aquatic Eco-Systems) using UV and solid filtration on a 14:10 L/D cycle at a temperature of 27°C. Fish were fed twice a day with Tetramin Tropical Flakes (Tetra, USA).

### Alarm Substance

We created a single batch of alarm substance following modified guidelines using 20 randomly selected donor fish (Speedie & Gerlai, 2008). In brief, donor fish were euthanized by rapid chilling followed by light abrasion of lateral skin cells on one side of each donor fish, ensuring that no blood was drawn. Donor bodies were then individually soaked in 10 mL of DI water for 10 minutes. We determined a working concentration through a pilot dose-response study (DI water, 10%, 50%, and 100% alarm substance). The 50% concentration elicited a significantly higher increase in freezing behavior compared to the DI water (*t*(22)= 3.24, *p* = .004, d = 2.33) and 10% (*t*(22)= 3.15, *p* = .005, d = 2.14) alarm substance administrations (Figure S1). We therefore selected 50% as the working concentration. A total of 200 mL was filtered, diluted in half, and stored in aliquots at −20° C until use.

### Contextual Fear Learning

To assess learning and memory we developed a novel contextual fear conditioning paradigm. Zebrafish were tested individually in an acrylic testing arena (16 x 16 x 10 cm) filled with 1.4 L of system water. The arenas were surrounded by opaque white plastic on the bottom and sides to serve as the contextual stimulus. A second context consisted of red plastic on the bottom with a picture of underwater plants on the side walls.

The paradigm consisted of three phases across 7 days of testing (Figure S2): acclimation, training, recall. Three days prior to testing, test subjects were moved from group housing into individual housing. On day one (acclimation phase), fish were individually placed in the testing arena to acclimate for 15 minutes and then returned to their home tank. Two hours later this was repeated in the second context. On day two (training phase), fish were trained to associate the white context with exposure to alarm substance over four learning trials. Each learning trial was 15 minutes long and was divided into three subsections. Fish acclimated to the chamber for the first five minutes, followed by five minutes of recording the conditioned fear response. After these 10 minutes, 1 mL of alarm substance was administered into the water through plastic tubing that came from outside of the testing arena. Following alarm substance exposure, the unconditioned fear response was recorded for five minutes. This was repeated for a total of four trials with 30 minutes between each. Between trials, we placed fish back into their individual housing, rinsed out the testing arenas, and refilled with 1.4 L of fresh system water. On days three and seven (recall phase), animals were re-exposed to both the neutral context and the conditioned context for 15 minutes each, with two hours between tests. For acclimation and recall testing, the order of context exposure was counterbalanced across individuals. All testing procedures were approved by the Institutional Animal Care and Use Committee of University of Nebraska at Omaha/University of Nebraska Medical Center (17-070-00-FC, 17-064-08-FC).

### Behavior Analysis

All trials were video-recorded from above and later analyzed with Noldus EthoVision XT (Noldus XT, Wageningen, Netherlands). For each trial, we quantified two measures as indicators of a conditioned response: freezing time and erratic movements. The subject was considered frozen if it moved less than 0.5 cm/s. Erratic movement duration was quantified using Ethovision’s Activity State analysis option (Noldus XT, Wageningen, Netherlands). The activity threshold was set to 99% and bins less than 0.1 seconds were removed. As erratic movements and freezing cannot occur simultaneously, we report duration of erratic movements as a proportion of total time spent moving. To validate software quantification of erratic movement duration, two independent observers manually recorded the duration of erratic movements for all of the unconditioned responses of the alarm substance group. Computer analyzed erratic movements were highly correlated with both observers (*r*_observer 1_ = 0.87, *p*_observer 1_ = 1.93*10^-15^ and *r*_observer 2_ = 0.91, *p*_observer 2_ = 2.77*10^-19^).

### Statistics

All statistics were performed using SPSS software (Version 24). To analyze freezing and erratic movement durations, we used three-way analysis of variance (ANOVA) models with strain, sex, and treatment group as between-subject factors. For analysis of acclimation on day one and memory recall at days three and seven, we used a repeated-measures three-way ANOVA with conditioned vs. neutral context as the within subjects factor. For analysis of the learning phase, we used a repeated-measures three-way ANOVA with the four conditioned response trials as the within-subjects factor. Individual comparisons were made with independent samples t-tests. Given the documented relationship between body size and boldness, we attempted to control for this by entering standard length into the models as a covariate (Brown & Braithwaite, 2004; Harris et al., 2010; Kern et al., 2016; Roy & Bhat, 2018). To account for multiple comparisons, we applied the Benjamini-Hochberg correction to determine significance (Benjamini et al., 2001). For all significant differences (p < 0.05) we also report the effect sizes (Cohen’s d (d) for t-tests and partial eta-squared (ηp^2^) for ANOVAs)(Wassertheil & Cohen, 1970). All effect sizes were medium or large effects (Richardson, 2011; Starkings, 2012; Wassertheil & Cohen, 1970).

## Results

During Day 1 acclimation there were no significant within-subjects effects of context or any interaction effect on baseline freezing or erratic movement behaviors. HSB fish froze significantly more than LSB fish overall (*F*_1, 55_ =10.81, *p* = .002, ηp^2^ = .16). However, there were no other significant between-subjects effects or interaction effects for freezing, nor any for erratic movements (all *p* > .05; Figure S3)

During the training phase (Day 2), fish that received alarm substance showed a significantly higher unconditioned response for freezing (*F*_1, 55_ = 563.41, *p* = 1.41*10^-30^, ηp^2^= .91) and erratic movements (*F*_1, 55_ = 11.77, *p* = .001, ηp^2^= .18) compared to DI water (Figure S4). There were no other significant between-subjects effects or interaction effects for the unconditioned fear response (all *p* > .05). In the conditioned fear response period, there was a significant trial*treatment group interaction effect for both freezing (*F*_3, 165_ = 71.31, *p* = 1.26*10^- 29^, ηp^2^= .57) and erratic movements (*F*_3, 165_ = 2.74, *p* = .045, ηp^2^= .05). The alarm substance group increased freezing across the four trials at a faster rate than the DI control group (Figure 1). For freezing behavior, there was a significant trial*strain*treatment group interaction (*F*_3, 165_ = 3.52, *p* = .016, ηp^2^= .06) where treated HSB fish increased freezing behavior at a faster rate than LSB fish. HSB fish exposed to alarm substance froze significantly more than LSB fish at trial two (*t*(46) = 3.29, *p* = .002, d = .95) and was not significant at trials one (*t*(46) = 1.78, *p* = .082), three (*t*(46) = 1.97, *p* = .055), or four (*t*(46) = 1.33, p = .189). Full model results are presented in Table S2.

**Figure 1.**
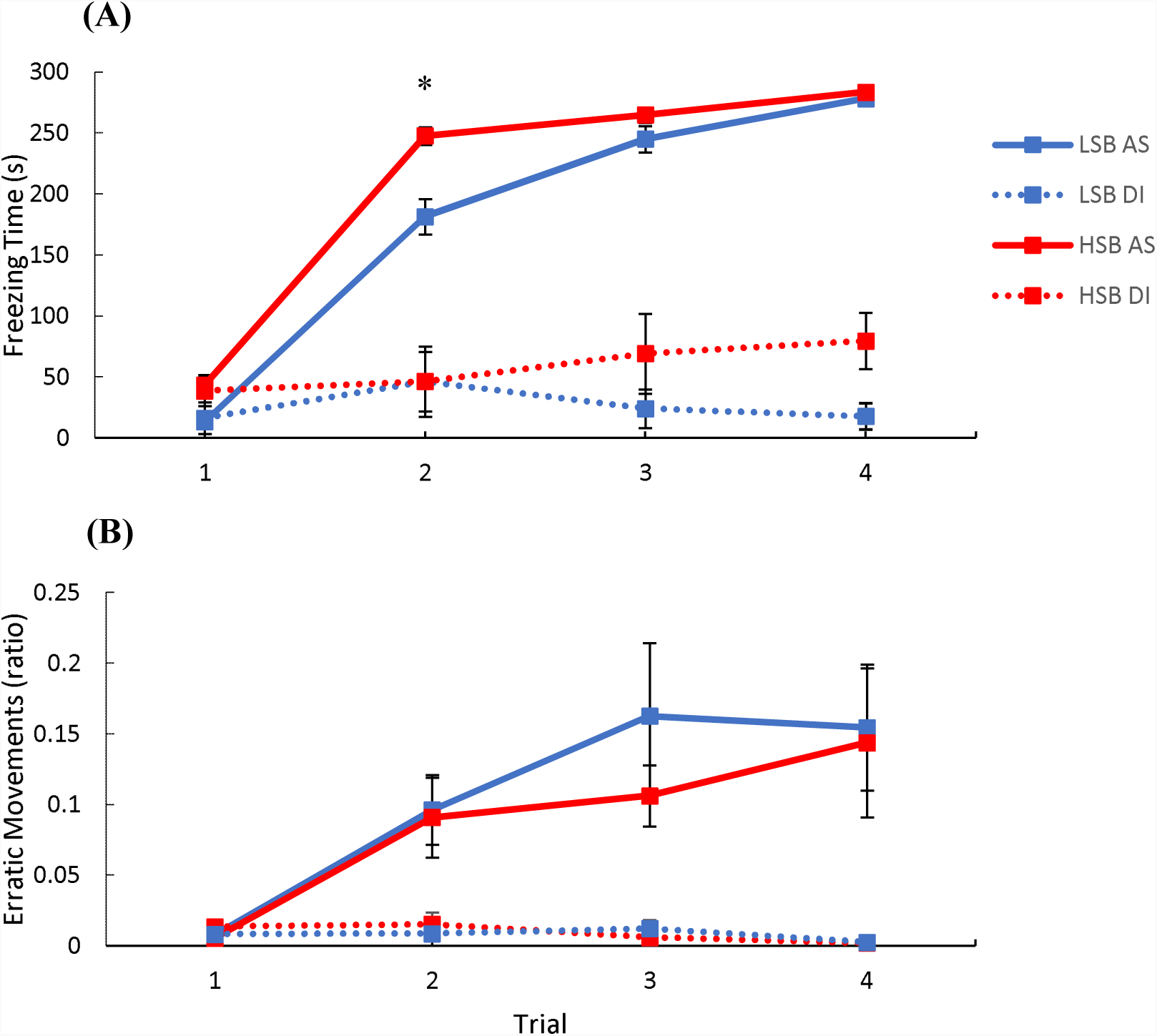
Acquisition of fear memory over four training trials. Freezing time (A) and erratic movement ratio (B) were measured for high stationary behavior (HSB) and low stationary behavior (LSB) fish exposed to distilled water (DI) or alarm substance (AS). Points represent mean ± 1 standard error. * indicates *p* < .05 for within-treatment group comparison.

During memory recall testing there was a significant context*treatment group interaction effect for both behaviors at 24h (Freezing: *F*_1, 55_ = 49.45, *p* = 2.97*10^-9^, ηp^2^= .48, erratic movements: *F*_1, 55_ = 5.41, *p* = .024, ηp^2^= .09, Figure 2) and freezing behavior at 96h (*F*_1, 55_ = 8.03, *p* = .006, ηp^2^= .127, Figure 3) post-training. In the alarm substance, but not the DI water group, both strains displayed significantly higher antipredatory behaviors in the conditioned context compared to the neutral context. At 96 hours post-training, there was a significant strain*treatment interaction effect for freezing behavior (*F*_1, 55_ = 4.13, *p* = .047, ηp^2^= .07). Treated HSB fish showed significantly higher freezing behavior compared to treated LSB fish in the conditioned context at 96h (*t*(46) = 3.62, *p* = .001, d = 1.01). Full model results are presented in Table S2.

**Figure 2.**
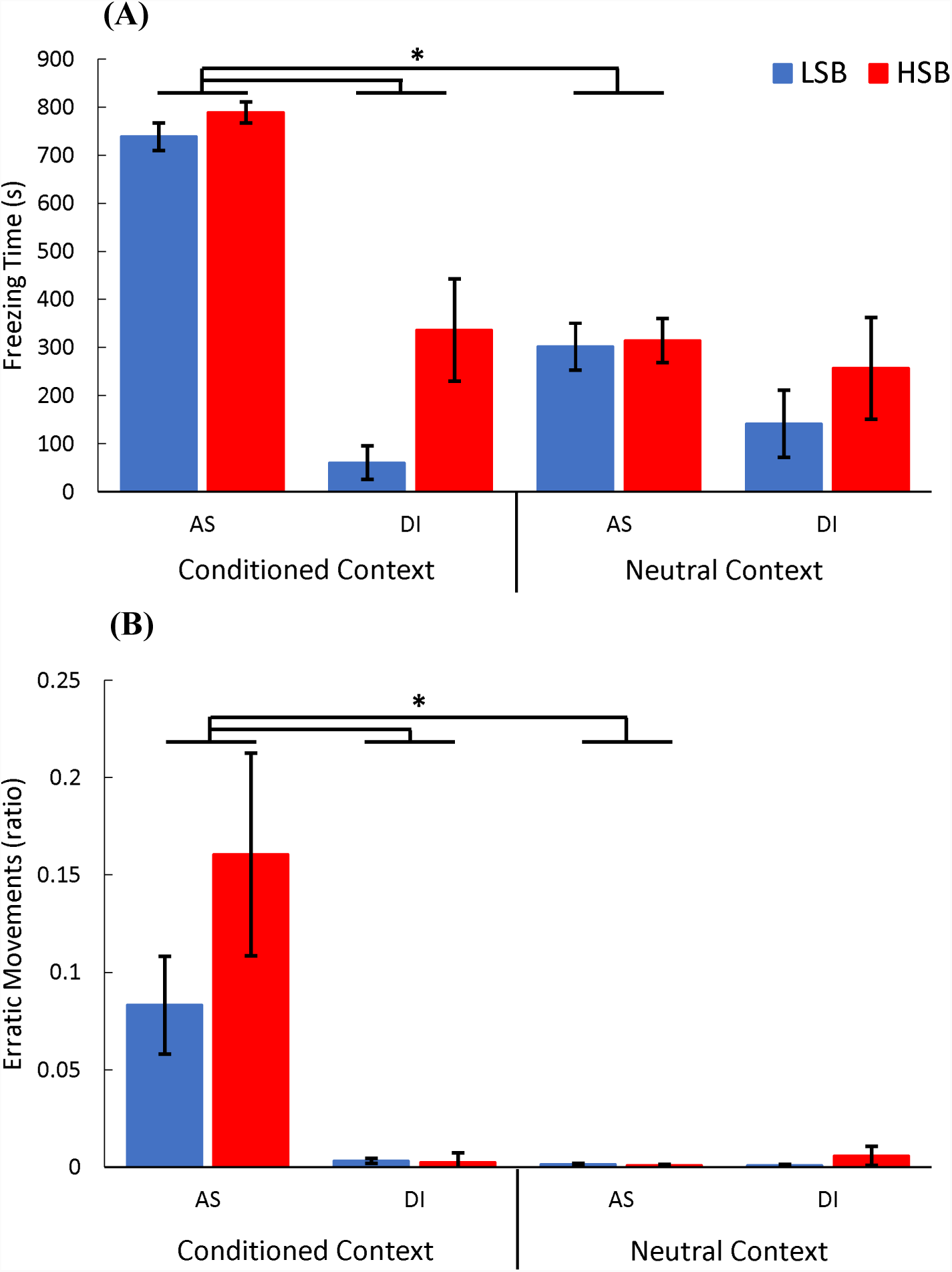
Fear memory recall 24 hours post-training. We measured freezing time (A) and erratic movement ratio (B) for high stationary behavior (HSB) and low stationary behavior (LSB) fish exposed to distilled water (DI) or alarm substance (AS) during training. Bars represent mean ± 1 standard error in the conditioned context and neutral context. * indicates *p* < .05.

**Figure 3.**
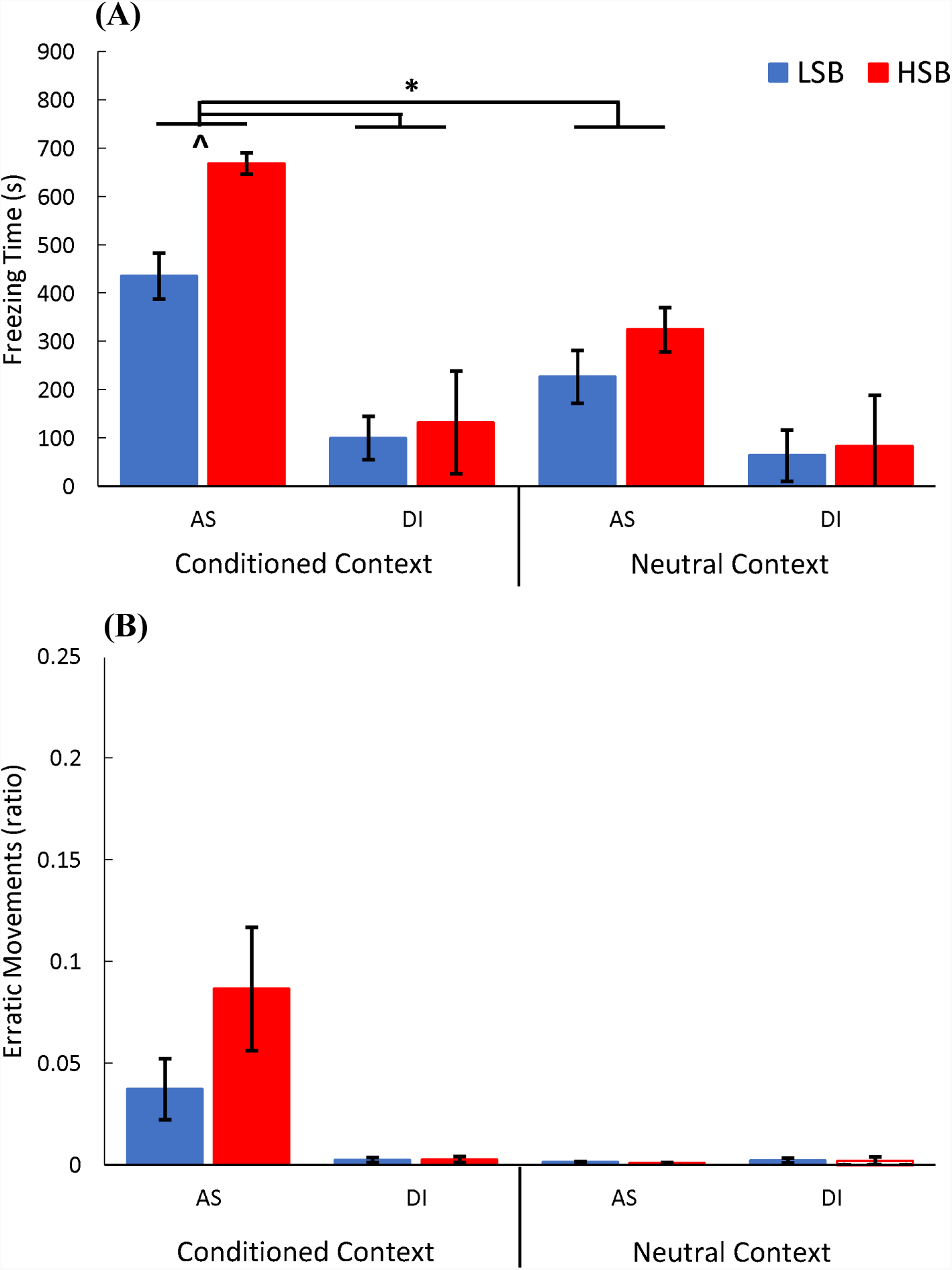
Fear memory recall 96 hours post-training. We measured freezing time (A) and erratic movement ratio (B) for high stationary behavior (HSB) and low stationary behavior (LSB) fish exposed to distilled water (DI) or alarm substance (AS) during training. Bars represent mean ± 1 standard error in the conditioned context and neutral context. * indicates *p* < .05. ^ indicates p <.05 for within-treatment group comparison in the conditioned context.

## Discussion

While it is essential for animals to encode and recall salient experiences, it is unclear how different stress coping strategies may influence the use of contextual information to predict and avoid danger in the future. In the present study, we measured the learning rate and duration of a fear memory in selectively-bred lines of zebrafish that display proactive and reactive coping styles. Overall, we found that reactive zebrafish more readily associated a fearful olfactory stimulus with contextual information and retained this fear memory longer compared to proactive individuals. We did not observe any sex differences in contextual fear learning or memory.

Learning rate and memory duration can differ amongst individuals with different personality types (Lucon-Xiccato & Bisazza, 2017; Sih & Del Giudice, 2012). We observed that reactive zebrafish (HSB strain) acquire a contextual fear memory at a significantly faster rate than proactive zebrafish (LSB strain) (Figure 2). With higher tendencies to exhibit risk-averse behaviors and elevated cortisol responses, reactive individuals may perceive stressors as more threatening, which could facilitate faster encoding of aversive experiences. Faster learning rates in reactive individuals have also been observed in other teleost (Budaev & Zhuikov, 1998; Mesquita et al., 2015) and avian species (Exnerová et al., 2010; Miller et al., 2006). While studies have documented faster learning proactive individuals (Amy et al., 2012; Bolhuis et al., 2004; DePasquale et al., 2014; Dugatkin & Alfieri, 2003; Mazza et al., 2018; Trompf & Brown, 2014), this may be due to different learning tasks or type of reinforcing stimulus. Reactive individuals have higher learning performance with aversive conditioning whereas proactive individuals tend to learn more quickly in exploratory or discrimination tasks with appetitive conditioning (Bolhuis et al., 2004; Budaev & Zhuikov, 1998; DePasquale et al., 2014; Dugatkin & Alfieri, 2003; Mesquita et al., 2015). It is unlikely innate contextual preferences could explain our results as there was no significant difference in freezing during acclimation between the conditioned or neutral context for either strains (Figure S3). Similarly, with no significant strain differences in freezing and erratic behaviors after first exposure to the alarm substance (unconditioned fear response period during first learning trial), it is also unlikely the strains have different response thresholds (Figure S4).

Freezing time and erratic movements during the recall phase indicated that both strains recalled the fear memory at least four days following training. However, the HSB fish showed significantly higher levels of freezing in the conditioned context at 96 hours suggesting that reactive individuals encode a more resilient fear memory than proactive individuals (Figure 3). Differences in learning and memory between stress coping styles are seen in both contextual (e.g. general environment) and cued (e.g. specific neutral odors or visual stimuli) learning of salient information using a threatening stimulus. Animals displaying a reactive coping style may repress exploratory behavior and be more risk-averse for longer when re-exposed to potentially dangerous contexts or cues to minimize risks of injury. This interpretation is consistent with other studies suggesting that reactive individuals retain fearful memories for longer (Brown et al., 2013; Exnerová et al., 2010). However, one study found that proactive rainbow trout retained a conditioned fear response for longer, which may be due to the reactive trout having faster extinction learning (Moreira et al., 2004). We speculate that differences in the rate of formation and duration of associations between aversive stimuli and an environmental context (e.g. microhabitat) may shape subsequent resource utilization (e.g. alter foraging routes, exploration range, duration of behavioral displays) resulting in altered population dynamics and compositions in the wild. Studies show that predation levels in a given habitat can influence learning and memory behaviors at the population level where individuals from low predation habitats tend to display higher activity and exploration (more proactive) and faster spatial learning capabilities to find food resources (Brown & Braithwaite, 2005; Brydges et al., 2008; DePasquale et al., 2014). While outside the scope of the current study, future studies should examine whether contextual learning under wild conditions alters habitat use and how it differs between individuals of alternative stress coping styles.

Painful or frightening stimuli can quickly modify current and future behavioral responses. Studies using electric shocks in fear conditioning have revealed important insights into the proximate mechanisms of learning and memory (Maren, 2001; Maren et al., 2013). However, electric shocks have limited ecological relevance to the evolution of adaptive animal behavior. Predator odors or chemical alarm signals are alternative, but ecologically relevant aversive conditioning stimuli. While alarm substance is used as an aversive conditioning stimulus in other studies utilizing teleosts (Brosnan et al., 2003; Brown et al., 2013; Hall & Suboski, 1995; Ruhl et al., 2017), our conditioning paradigm allows for effective analysis of behavior at the individual level and achieved an unconditioned response rate in ∼80% of fish. Further, alarm substance induced similar unconditioned fear responses in all fish (Figure S4). Only fish exposed to alarm substance displayed increasing conditioned fear responses across learning trials (Figure 1) and had higher levels in the conditioned context during memory recall (Figures 2, 3). This is consistent with freezing and avoidance behaviors observed in other fear conditioning paradigms utilizing chemical alarm signals and electric shocks (Brown et al., 2011; Kenney et al., 2017; Takahashi et al., 2008). Collectively this suggests that all fish acquired the association between the alarm substance and the contextual information, and were able to discriminate between the conditioned and neutral contexts. Further, freezing behavior shows strong consistent individual differences and is highly repeatable in both of the proactive and reactive zebrafish strains used in this study (Baker et al., 2018). Ecologically-relevant stimuli like alarm substance may help elucidate adaptive cognitive processes in response to predation or other selective pressures (Kim & Jung, 2018; Pellman & Kim, 2016).

Differences in cognition between proactive and reactive stress coping styles are observed across various taxonomic groups, which suggest common underlying neuromolecular mechanisms. Interestingly, key mechanisms for learning and memory (neural plasticity and neurogenesis), are elevated in reactive individuals which could bias learning and memory capabilities (Øverli & Sørensen, 2016; Sørensen et al., 2013; Wong et al., 2015). Additionally, variation in cognitive flexibility among stress coping styles has been linked to key neurotransmitter systems (e.g. dopaminergic, serotonergic, GABAergic)(Banuelos et al., 2014; Beas et al., 2016; Coppens et al., 2010; Höglund et al., 2017; Wong et al., 2015). Consistent with this idea, basal expression of genes in the brain related to neural plasticity and neurotransmission are differentially regulated between the HSB and LSB strains (Wong et al., 2015). We hypothesize that faster fear learning rates and stronger memory recall of reactive individuals in this study is facilitated by altered expression of these genes in response to fearful stimuli. Selectively bred proactive and reactive behavioral phenotypes will be useful in investigating these proximate mechanisms of cognitive biases and other correlated traits in future studies.

## Conclusion

Intriguingly we document several interaction effects between an individual’s stress coping style and learning and memory of a fearful association. Specifically, despite showing similar acute responses to potential predation, we find that reactive individuals actively encode this information more quickly and that it lasts longer than proactive individuals. Alternatively, proactive individuals may forget or suppress fearful associations sooner to maximize future resource acquisition. We also show that alarm substance can be used to understand contextual learning and memory differences between stress coping styles (i.e. personality types). It is important to consider a variety of paradigms as different associations and reinforcement valences may incur different sets of tradeoffs that influence cognition. Lastly, these behavioral findings present a promising basis to investigate the neuromolecular mechanisms underlying cognitive biases and stress coping styles.

## Acknowledgements

We are grateful to D. Revers and A. Park for zebrafish husbandry. We thank A. Goodman and S. Bresnahan for helpful discussions and S. Bresnahan for assisting in manual scoring of erratic movements.

## Supplementary Information

**Table S1.** Results of repeated measures GLM for the acquisition learning phase for freezing time and erratic movement ratio.

|  | Freezing Time | Erratic Movement |
| --- | --- | --- |
| | $F_{(p, \eta p^2)}$ | $F_{(p, \eta p^2)}$ |
| Within-Subjects Effects ($df = 3, 165$) | | |
| Trial | **3.42** (.019, .06) | 1.35 (.261) |
| Trial*Strain | 0.18 (.194) | 0.21 (.892) |
| Trial*Sex | 1.18 (.318) | 0.80 (.496) |
| Trial*Treatment | **71.31** ($1.26 \times 10^{-29}$, .57) | **2.74** (.045, .05) |
| Trial*Strain*Sex | 1.49 (.220) | 0.69 (.560) |
| Trial*Strain*Treatment | **3.52** (.016, .06) | 0.16 (.921) |
| Trial*Sex*Treatment | 1.29 (.281) | 0.48 (.696) |
| Trial*Strain*Sex*Treatment | 0.45 (.720) | 0.63 (.600) |
| Between Subjects Effects ($df = 1, 55$) | | |
| Intercept | **7.63** (.008, .12) | 2.50 (.120) |
| Strain | **13.20** (.001, .19) | 0.11 (.740) |
| Sex | **14.01** ($4.36 \times 10^{-4}$, .20) | 0.62 (.433) |
| Treatment | **375.76** ($3.00 \times 10^{-26}$, .87) | **16.25** ($1.72 \times 10^{-4}$, .23) |
| Strain*Sex | 0.48 (.490) | 1.37 (.247) |
| Strain*Treatment | 0.08 (.783) | 0.01 (.918) |
| Sex*Treatment | **10.27** (.002, .16) | 0.01 (.937) |
| Strain*Sex*Treatment | 3.42 (.070) | 0.93 (.338) |

Bold text indicates $p < 0.05$

**Table S2.** Results of repeated measures GLM for the memory recall phase for freezing time and erratic movement ratio at 24h and 96h post training.

|  | 24h Freezing Time | 24h Erratic Movement | 96h Freezing Time | 96h Erratic Movement |
| --- | --- | --- | --- | --- |
| | $F_{(p, \eta p^2)}$ | $F_{(p, \eta p^2)}$ | $F_{(p, \eta p^2)}$ | $F_{(p, \eta p^2)}$ |
| Within-Subjects Effects (df = 1, 55) |  |  |  |  |
| Context | 1.21 (.277) | 0.02 (.900) | 3.31 (.074) | 0.10 (.755) |
| Context*Strain | 1.82 (.518) | 0.63 (.430) | 0.10 (.754) | 0.79 (.378) |
| Context*Sex | 0.06 (.805) | 0.09 (.762) | 0.94 (.336) | 0.42 (.521) |
| Context*Treatment | **49.45** ($2.97 \times 10^{-9}$, .48) | **5.41** (.024, .09) | **8.03** (.007, .13) | 3.54 (.065) |
| Context*Strain*Sex | 0.83 (.365) | 0.02 (.900) | 0.82 (.370) | 0.00 (.963) |
| Context*Strain*Treatment | 1.04 (.312) | 0.82 (.369) | 0.12 (.726) | 0.68 (.413) |
| Context*Sex*Treatment | 0.89 (.351) | 0.05 (.823) | 1.79 (.187) | 0.67 (.415) |
| Context*Strain*Sex*Treatment | 0.52 (.472) | 0.01 (.946) | 0.22 (.645) | 0.03 (.862) |
| Between Subjects Effects (df = 1, 55) |  |  |  |  |
| Intercept | 0.07 (.791) | 0.01 (.928) | 0.32 (.572) | 0.12 (.735) |
| Strain | 7.17* (.010, .03) | 0.78 (.382) | **7.60** (.009, .12) | 0.24 (.630) |
| Sex | 0.49 (.488) | 0.06 (.802) | 0.26 (.613) | 0.76 (.387) |
| Treatment | **51.31** ($1.97 \times 10^{-9}$, .483) | **4.99** (.030, .08) | **51.15** ($2.75 \times 10^{-9}$, .48) | 3.27 (.076) |
| Strain*Sex | 1.52 (.223) | 0.00 (.998) | 1.77 (.188) | 0.02 (.899) |
| Strain*Treatment | 3.47 (.068) | 0.65 (.425) | **4.13** (.047, .07) | 0.65 (.423) |
| Sex*Treatment | **6.65** (.013, .11) | 0.12 (.727) | 1.11 (.296) | 0.65 (.424) |
| Strain*Sex*Treatment | **4.33** (.045, .07) | 0.04 (.845) | 0.01 (.909) | 0.00 (.989) |

Bold text indicates $p < 0.05$

**Figure S1.**
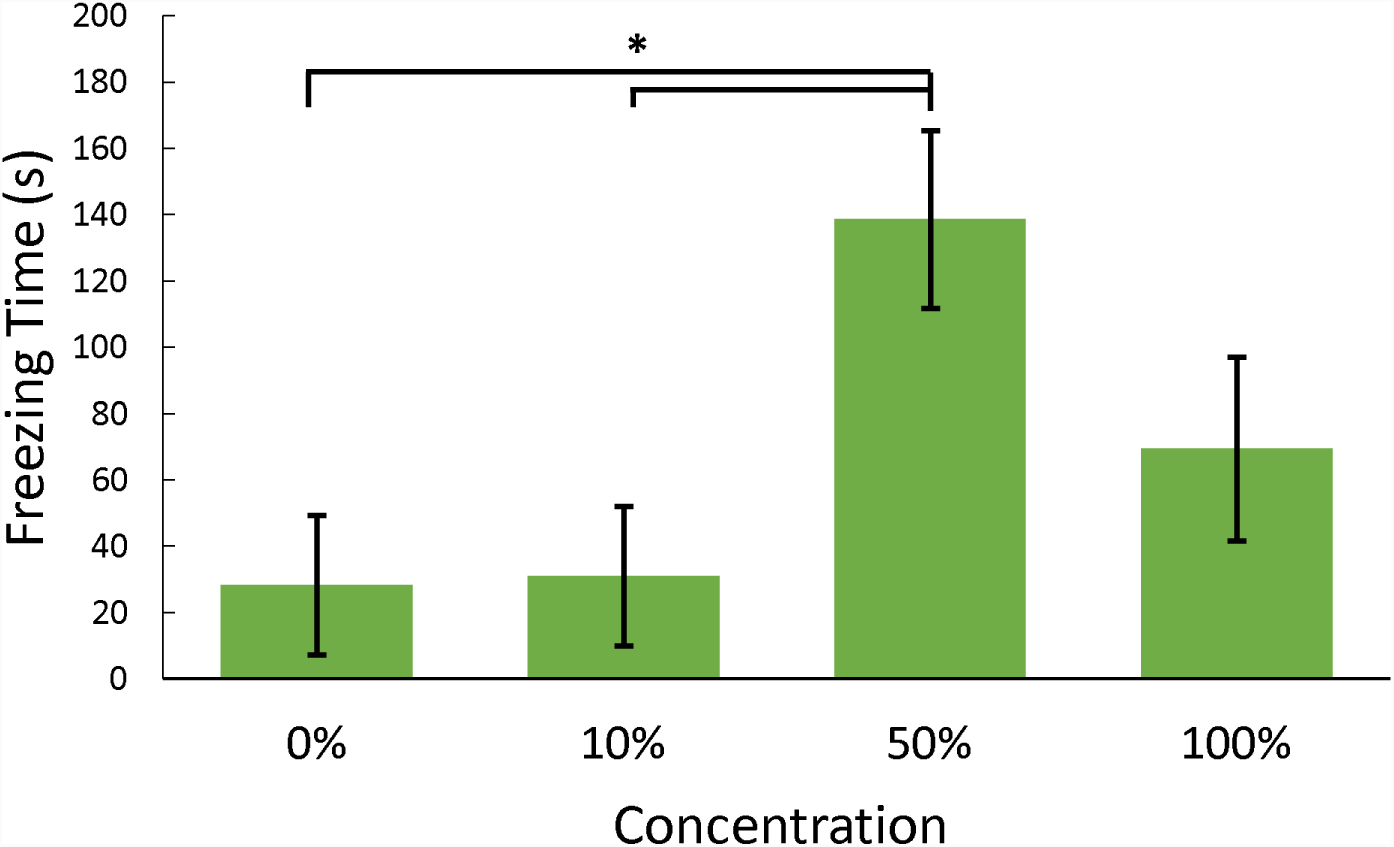
Dose response analysis of alarm substance administration on freezing behavior. For pilot trials, fish were recorded for five minutes after administration of four concentrations of alarm substance (DI water, 10%, 50%, 100%). Bars indicate mean ± 1 standard error. * indicates *p* < .05.

**Figure S2.**
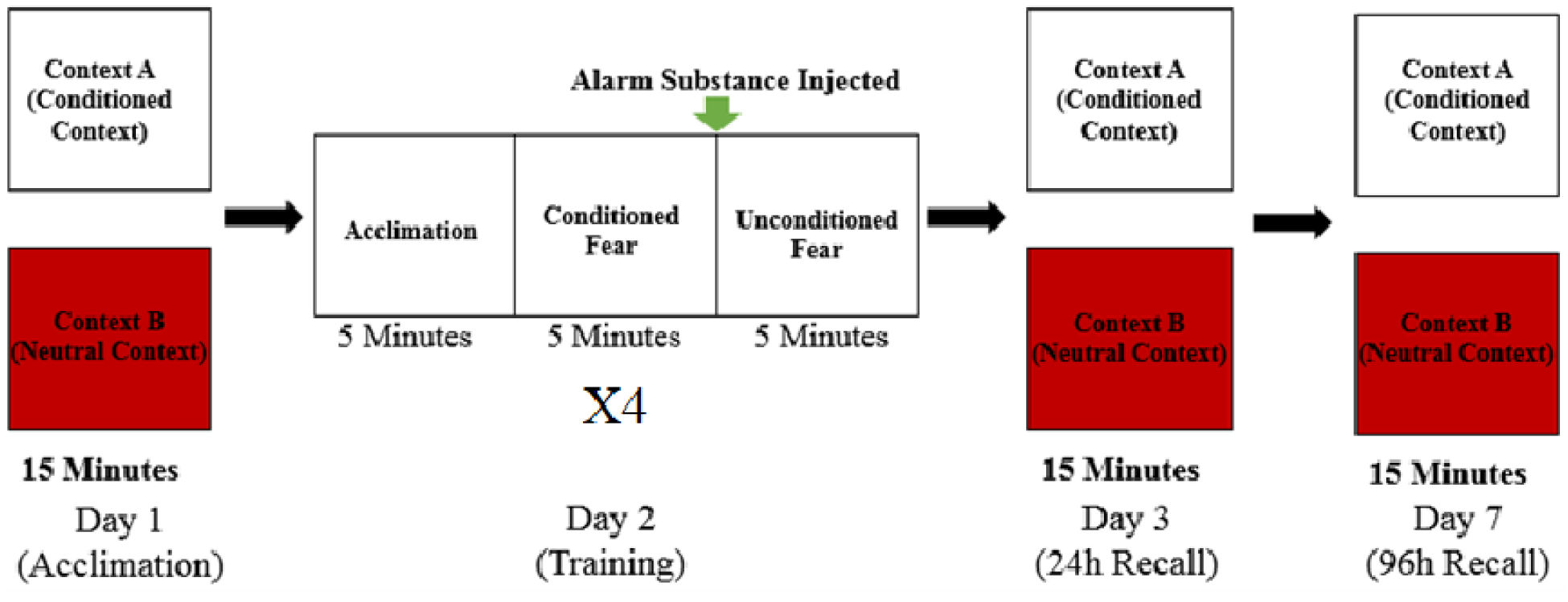
Contextual fear conditioning protocol. On day one, animals were exposed to both the conditioned and neutral contexts for 15 minutes to acclimate. On day two, fish were trained to associate alarm substance exposure to the conditioned context. Training trials consisted of three five minute blocks. For the first five minutes animals were allowed to acclimate to the arena. The second five minutes were recorded as an indicator of conditioned fear, and used to measure learning rate over four trials. Alarm substance was administered at the end of the conditioned fear block, and the fish’s unconditioned fear response was measured for five minutes. The training trial was repeated four times with 30 minutes in their home tank between trials. On days three and seven, memory recall was tested by re-exposing fish to the conditioned and neutral contexts for 15 minutes each with two hours between contexts.

**Figure S3.**
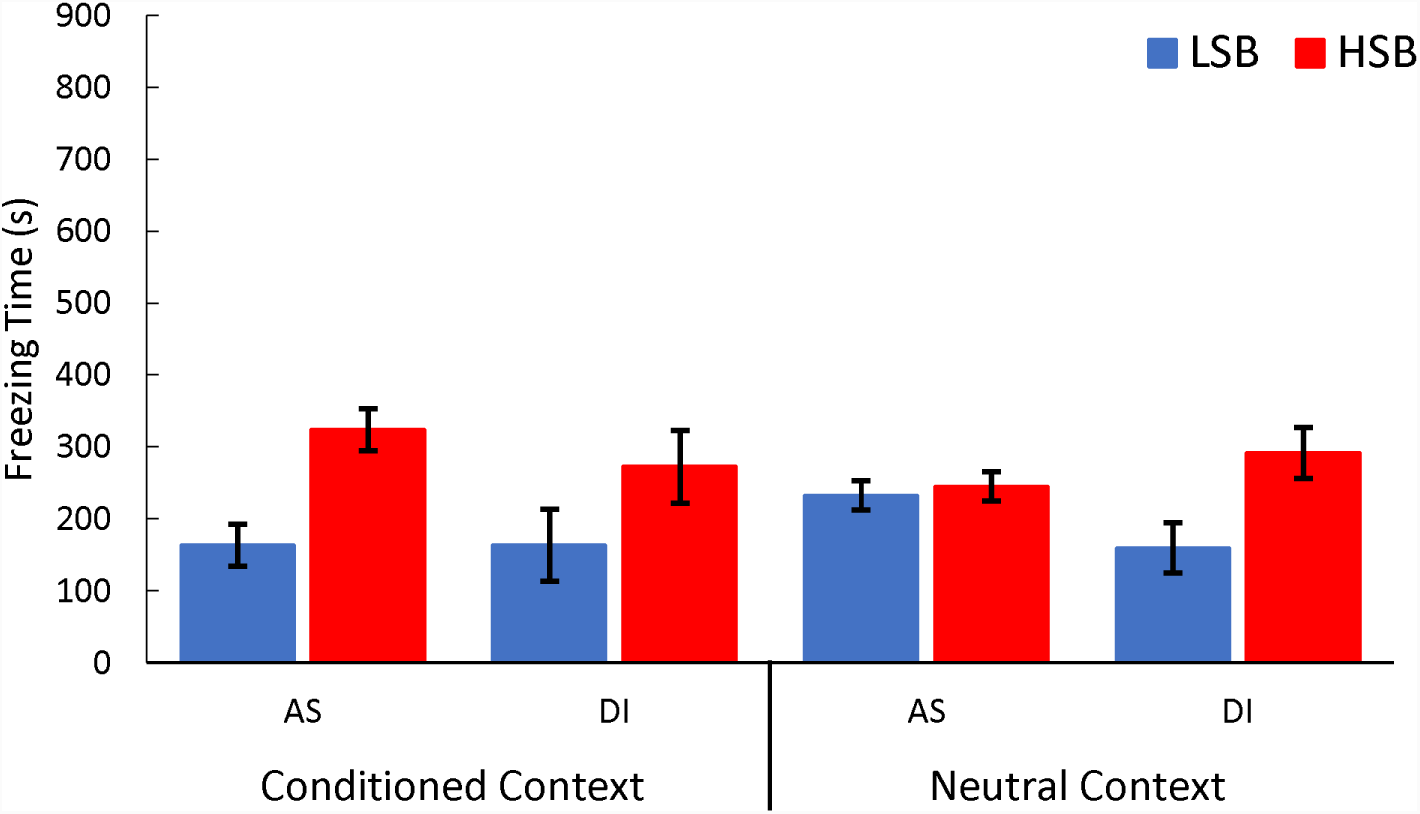
Freezing time displayed during acclimation phase. We measured freezing time for high stationary behavior (HSB) and low stationary behavior (LSB) fish exposed to distilled water (DI) or alarm substance (AS). Bars represent mean ± 1 standard error in the conditioned context and neutral context. Overall, HSB fish froze significantly more than LSB fish. However, there was no effect of context or treatment group on freezing time.

**Figure S4.**
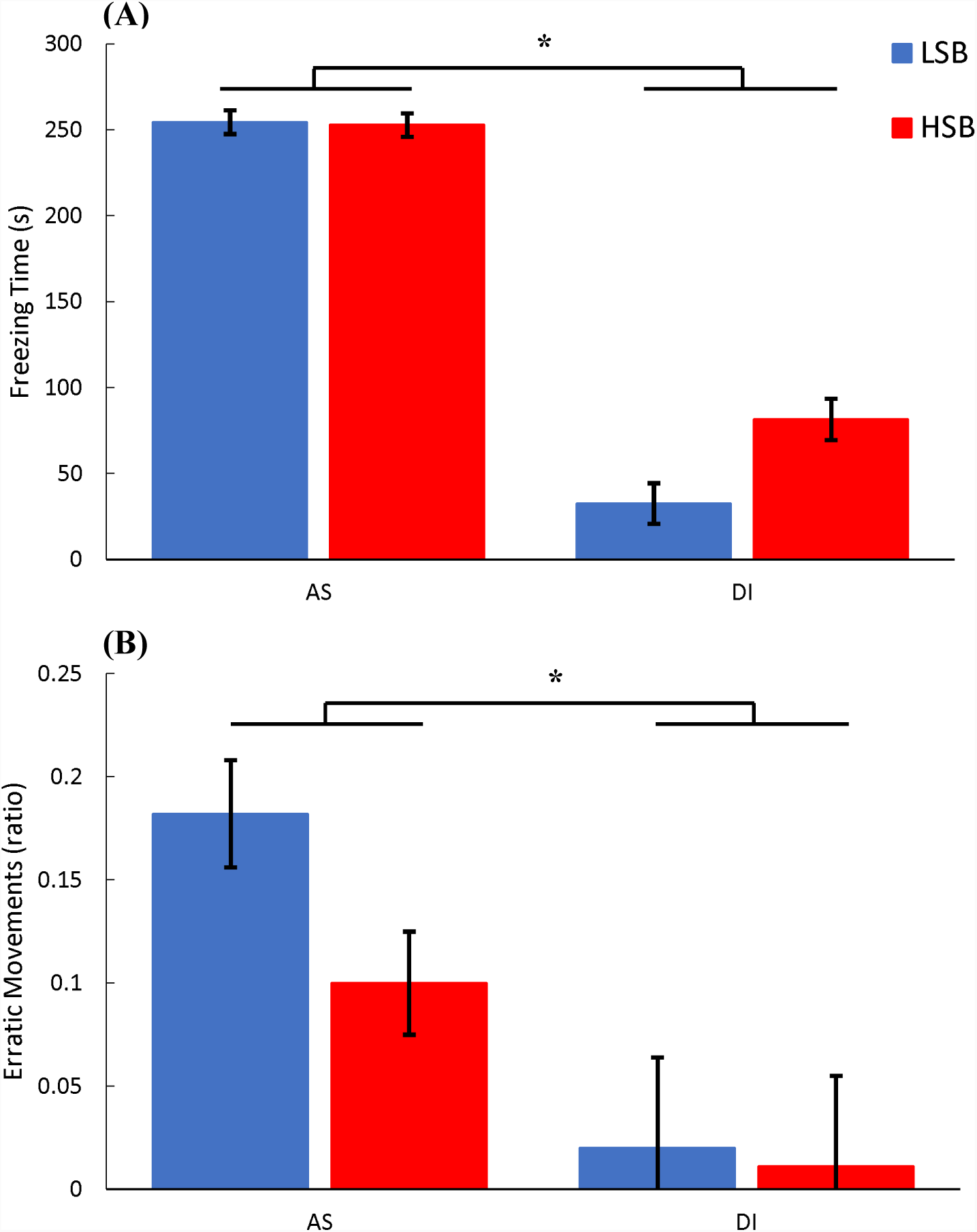
Unconditioned fear response during the first learning trial. We measured freezing time (A) and erratic movement ratio (B) for high stationary behavior (HSB) and low stationary behavior (LSB) fish exposed to distilled water (DI) or alarm substance (AS). Bars represent mean ± 1 standard error in the conditioned context. * indicates *p* < .05.

## References

Amy, M., van Oers, K., & Naguib, M. (2012). Worms under cover: Relationships between performance in learning tasks and personality in great tits (Parus major). Animal Cognition. https://doi.org/10.1007/s10071-012-0500-3

Baker, M. R., Goodman, A. C., Santo, J. B., & Wong, R. Y. (2018). Repeatability and reliability of exploratory behavior in proactive and reactive zebrafish, Danio rerio. Scientific Reports. https://doi.org/10.1038/s41598-018-30630-3

Baker, M. R., Hofmann, H. A., & Wong, R. Y. (n.d.). Neurogenomics of Behavioural Plasticity in Socioecological Contexts. https://doi.org/10.1002/9780470015902.a0026839

Banuelos, C., Beas, B. S., McQuail, J. A., Gilbert, R. J., Frazier, C. J., Setlow, B., & Bizon, J. L. (2014). Prefrontal Cortical GABAergic Dysfunction Contributes to Age-Related Working Memory Impairment. Journal of Neuroscience. https://doi.org/10.1523/JNEUROSCI.5192-13.2014

Beas, B. S., Setlow, B., & Bizon, J. L. (2016). Effects of acute administration of the GABA(B) receptor agonist baclofen on behavioral flexibility in rats. Psychopharmacology. https://doi.org/10.1007/s00213-016-4321-y

Benjamini, Y., Drai, D., Elmer, G., Kafkafi, N., & Golani, I. (2001). Controlling the false discovery rate in behavior genetics research. Behavioural Brain Research, 125(1–2), 279–284. https://doi.org/10.1016/S0166-4328(01)00297-2

Bolhuis, J. E., Schouten, W. G. P., Leeuw, J. A. De, Schrama, J. W., & Wiegant, V. M. (2004). Individual coping characteristics, rearing conditions and behavioural flexibility in pigs. Behavioural Brain Research, 152(2), 351–360. https://doi.org/10.1016/j.bbr.2003.10.024

Brosnan, S. F., Earley, R. L., & Dugatkin, L. A. (2003). Observational Learning and Predator Inspection in Guppies (Poecilia reticulata). Ethology. https://doi.org/10.1046/j.0179-1613.2003.00928.x

Brown, C., & Braithwaite, V. A. (2004). Size matters: A test of boldness in eight populations of the poeciliid Brachyraphis episcopi. Animal Behaviour. https://doi.org/10.1016/j.anbehav.2004.04.004

Brown, C., & Braithwaite, V. A. (2005). Effects of predation pressure on the cognitive ability of the poeciliid Brachyraphis episcopi. Behavioral Ecology. https://doi.org/10.1093/beheco/ari016

Brown, G. E., Ferrari, M. C. O., & Chivers, D. P. (2011). Learning about Danger: Chemical Alarm Cues and Threat-Sensitive Assessment of Predation Risk by Fishes. In Fish Cognition and Behavior. https://doi.org/10.1002/9781444342536.ch4

Brown, G. E., Ferrari, M. C. O., Malka, P. H., Fregeau, L., Kayello, L., & Chivers, D. P. (2013). Retention of acquired predator recognition among shy versus bold juvenile rainbow trout. Behavioral Ecology and Sociobiology, 67(1), 43–51. https://doi.org/10.1007/s00265-012-1422-4

Brydges, N. M., Heathcote, R. J. P., & Braithwaite, V. A. (2008). Habitat stability and predation pressure influence learning and memory in populations of three-spined sticklebacks. Animal Behaviour. https://doi.org/10.1016/j.anbehav.2007.08.005

Bshary, R., & Brown, C. (2014). Fish cognition. Current Biology. https://doi.org/10.1016/j.cub.2014.08.043

Budaev, S. V., & Zhuikov, A. Y. (1998). Avoidance learning and “personality” in the guppy (Poecilia reticulata). Journal of Comparative Psychology, 112(1), 92–94. https://doi.org/10.1037/0735-7036.112.1.92

Carere, C., & Locurto, C. (2011). Interaction between animal personality and animal cognition. Current Zoology. https://doi.org/10.1093/czoolo/57.4.491

Coppens, C. M., De Boer, S. F., & Koolhaas, J. M. (2010). Coping styles and behavioural flexibility: Towards underlying mechanisms. Philosophical Transactions of the Royal Society B: Biological Sciences. https://doi.org/10.1098/rstb.2010.0217

Dalesman, S. (2018). Habitat and social context affect memory phenotype, exploration and covariance among these traits. Philosophical Transactions of the Royal Society of London. Series B, Biological Sciences. https://doi.org/10.1098/rstb.2017.0291

DePasquale, C., Wagner, T., Archard, G. A., Ferguson, B., & Braithwaite, V. A. (2014). Learning rate and temperament in a high predation risk environment. Oecologia, 176(3), 661–667. https://doi.org/10.1007/s00442-014-3099-z

Dougherty, L. R., & Guillette, L. M. (2018). Linking personality and cognition: a meta-analysis. Philosophical Transactions of the Royal Society B: Biological Sciences. https://doi.org/10.1098/rstb.2017.0282

Dugatkin, L. A., & Alfieri, M. S. (2003). Boldness, behavioral inhibition and learning. Ethology Ecology and Evolution, 15(1), 43–49. https://doi.org/10.1080/08927014.2003.9522689

Exnerová, A., Svádová, K. H., Fučíková, E., Drent, P., & Štys, P. (2010). Personality matters: Individual variation in reactions of naive bird predators to aposematic prey. Proceedings of the Royal Society B: Biological Sciences. https://doi.org/10.1098/rspb.2009.1673

Gaikwad, S., Stewart, A., Hart, P., Wong, K., Piet, V., Cachat, J., & Kalueff, A. V. (2011). Acute stress disrupts performance of zebrafish in the cued and spatial memory tests: The utility of fish models to study stress-memory interplay. Behavioural Processes. https://doi.org/10.1016/j.beproc.2011.04.004

Gerlai, R. (2010). Zebrafish antipredatory responses: A future for translational research? Behavioural Brain Research. https://doi.org/10.1016/j.bbr.2009.10.008

Gerlai, R. (2016). Learning and memory in zebrafish (Danio rerio). In Methods in Cell Biology. https://doi.org/10.1016/bs.mcb.2016.02.005

Griffin, A. S., Guillette, L. M., & Healy, S. D. (2015). Cognition and personality: An analysis of an emerging field. Trends in Ecology and Evolution. https://doi.org/10.1016/j.tree.2015.01.012

Hall, D., & Suboski, M. D. (1995). Visual and olfactory stimuli in learned release of alarm reactions by zebra danio fish (brachydanio rerio). Neurobiology of Learning and Memory, 63(3), 229–240. https://doi.org/10.1006/nlme.1995.1027

Harris, S., Ramnarine, I. W., Smith, H. G., & Pettersson, L. B. (2010). Picking personalities apart: Estimating the influence of predation, sex and body size on boldness in the guppy Poecilia reticulata. Oikos. https://doi.org/10.1111/j.1600-0706.2010.18028.x

Höglund, E., Silva, P. I. M., Vindas, M. A., & Øverli, Ø. (2017). Contrasting coping styles meet the wall: A dopamine driven dichotomy in behavior and cognition. Frontiers in Neuroscience. https://doi.org/10.3389/fnins.2017.00383

Ijaz, S., & Hoffman, E. J. (2016). Zebrafish: A Translational Model System for Studying Neuropsychiatric Disorders. Journal of the American Academy of Child and Adolescent Psychiatry, 55(9), 746–748. https://doi.org/10.1016/j.jaac.2016.06.008

Kenney, J. W., Scott, I. C., Josselyn, S. A., & Frankland, P. W. (2017). Contextual fear conditioning in zebrafish. Learning & Memory, 24(10), 516–523. https://doi.org/10.1101/lm.045690.117

Kern, E. M. A., Robinson, D., Gass, E., Godwin, J., & Langerhans, R. B. (2016). Correlated evolution of personality, morphology and performance. Animal Behaviour, 117, 79–86. https://doi.org/10.1016/j.anbehav.2016.04.007

Kim, J. J., & Jung, M. W. (2018). Fear paradigms: The times they are a-changin’. Current Opinion in Behavioral Sciences. https://doi.org/10.1016/j.cobeha.2018.02.007

Koolhaas, J. M., de Boer, S. F., Coppens, C. M., & Buwalda, B. (2010). Neuroendocrinology of coping styles: Towards understanding the biology of individual variation. Frontiers in Neuroendocrinology. https://doi.org/10.1016/j.yfrne.2010.04.001

Koolhaas, J. M., Korte, S. M., De Boer, S. F., Van Der Vegt, B. J., Van Reenen, C. G., Hopster, H., … Blokhuis, H. J. (1999). Coping styles in animals: Current status in behavior and stress-physiology. Neuroscience and Biobehavioral Reviews, 23(7), 925–935. https://doi.org/10.1016/S0149-7634(99)00026-3

Lucon-Xiccato, T., & Bisazza, A. (2017). Individual differences in cognition among teleost fishes. Behavioural Processes. https://doi.org/10.1016/j.beproc.2017.01.015

Maren, S. (2001). Neurobiology of Pavlovian fear conditioning. Annual Review of Neuroscience. https://doi.org/10.1146/annurev.neuro.24.1.897

Maren, S., Phan, K. L., & Liberzon, I. (2013). The contextual brain: Implications for fear conditioning, extinction and psychopathology. Nature Reviews Neuroscience. https://doi.org/10.1038/nrn3492

Mazza, V., Eccard, J. A., Zaccaroni, M., Jacob, J., & Dammhahn, M. (2018). The fast and the flexible: cognitive style drives individual variation in cognition in a small mammal. Animal Behaviour. https://doi.org/10.1016/j.anbehav.2018.01.011

Mesquita, F. O., Borcato, F. L., & Huntingford, F. A. (2015). Cue-based and algorithmic learning in common carp: A possible link to stress coping style. Behavioural Processes, 115, 25–29. https://doi.org/10.1016/j.beproc.2015.02.017

Miller, K. A., Garner, J. P., & Mench, J. A. (2006). Is fearfulness a trait that can be measured with behavioural tests? A validation of four fear tests for Japanese quail. Animal Behaviour. https://doi.org/10.1016/j.anbehav.2005.08.018

Miller, N. (2017). Cognition in fishes. Behavioural Processes. https://doi.org/10.1016/j.beproc.2017.03.013

Moreira, P. S. A., Pulman, K. G. T., & Pottinger, T. G. (2004). Extinction of a Conditioned Response in Rainbow Trout Selected for High or Low Responsiveness to Stress. Hormones and Behavior. https://doi.org/10.1016/j.yhbeh.2004.05.003

Norton, W.. b, & Bally-Cuif, L.. b. (2010). Adult zebrafish as a model organism for behavioural genetics. BMC Neuroscience. https://doi.org/10.1186/1471-2202-11-90

Oliveira, R. F. (2013). Mind the fish: zebrafish as a model in cognitive social neuroscience. Frontiers in Neural Circuits. https://doi.org/10.3389/fncir.2013.00131

Oswald, M. E., Drew, R. E., Racine, M., Murdoch, G. K., & Robison, B. D. (2012). Is Behavioral Variation along the Bold-Shy Continuum Associated with Variation in the Stress Axis in Zebrafish? Physiological and Biochemical Zoology, 85(6), 718–728. https://doi.org/10.1086/668203

Oswald, M. E., Singer, M., & Robison, B. D. (2013). The Quantitative Genetic Architecture of the Bold-Shy Continuum in Zebrafish, Danio rerio. PLoS ONE, 8(7). https://doi.org/10.1371/journal.pone.0068828

Øverli, Ø., & Sørensen, C. (2016). On the role of neurogenesis and neural plasticity in the evolution of animal personalities and stress coping styles. Brain, Behavior and Evolution. https://doi.org/10.1159/000447085

Øverli, Ø., Sørensen, C., Pulman, K. G. T., Pottinger, T. G., Korzan, W., Summers, C. H., & Nilsson, G. E. (2007). Evolutionary background for stress-coping styles: Relationships between physiological, behavioral, and cognitive traits in non-mammalian vertebrates. Neuroscience and Biobehavioral Reviews. https://doi.org/10.1016/j.neubiorev.2006.10.006

Pellman, B. A., & Kim, J. J. (2016). What Can Ethobehavioral Studies Tell Us about the Brain’s Fear System? Trends in Neurosciences. https://doi.org/10.1016/j.tins.2016.04.001

Richardson, J. T. E. (2011). Eta squared and partial eta squared as measures of effect size in educational research. Educational Research Review. https://doi.org/10.1016/j.edurev.2010.12.001

Roy, T., & Bhat, A. (2018). Population, sex and body size: determinants of behavioural variations and behavioural correlations among wild zebrafish *Danio rerio*. Royal Society Open Science, 5(1), 170978. https://doi.org/10.1098/rsos.170978

Ruhl, T., Zeymer, M., & von der Emde, G. (2017). Cannabinoid modulation of zebrafish fear learning and its functional analysis investigated by c-Fos expression. Pharmacology Biochemistry and Behavior, 153, 18–31. https://doi.org/10.1016/j.pbb.2016.12.005

Russ, J. (2018). Differences in Behavioral and Physiological Responses to Stress in Zebrafish. University of Nebraska at Omaha.

Sih, A., Bell, A., & Johnson, J. C. (2004). Behavioral syndromes: An ecological and evolutionary overview. Trends in Ecology and Evolution. https://doi.org/10.1016/j.tree.2004.04.009

Sih, A., & Del Giudice, M. (2012). Linking behavioural syndromes and cognition: a behavioural ecology perspective. Philosophical Transactions of the Royal Society B: Biological Sciences, 367(1603), 2762–2772. https://doi.org/10.1098/rstb.2012.0216

Sorato, E., Zidar, J., Garnham, L., Wilson, A., & Løvlie, H. (2018). Heritabilities and co-variation among cognitive traits in red junglefowl. Philosophical Transactions of the Royal Society B: Biological Sciences. https://doi.org/10.1098/rstb.2017.0285

Sørensen, C., Johansen, I. B., & Øverli, Ø. (2013). Neural plasticity and stress coping in teleost fishes. General and Comparative Endocrinology. https://doi.org/10.1016/j.ygcen.2012.12.003

Speedie, N., & Gerlai, R. (2008). Alarm substance induced behavioral responses in zebrafish (Danio rerio). Behavioural Brain Research, 188(1), 168–177. https://doi.org/10.1016/j.bbr.2007.10.031

Starkings, S. (2012). IBM SPSS Statistics 19 Made Simple by Colin D. Gray and Paul R. Kinnear. International Statistical Review. https://doi.org/10.1111/j.1751-5823.2012.00187_13.x

Takahashi, L. K., Chan, M. M., & Pilar, M. L. (2008). Predator odor fear conditioning: Current perspectives and new directions. Neuroscience and Biobehavioral Reviews. https://doi.org/10.1016/j.neubiorev.2008.06.001

Trompf, L., & Brown, C. (2014). Personality affects learning and trade-offs between private and social information in guppies, poecilia reticulata. Animal Behaviour, 88, 99–106. https://doi.org/10.1016/j.anbehav.2013.11.022

Wassertheil, S., & Cohen, J. (1970). Statistical Power Analysis for the Behavioral Sciences. Biometrics. https://doi.org/10.2307/2529115

Wong, R. Y., & Godwin, J. (2015). Neurotranscriptome profiles of multiple zebrafish strains. Genomics Data, 5, 206–209. https://doi.org/10.1016/j.gdata.2015.06.004

Wong, R. Y., Lamm, M. S., & Godwin, J. (2015). Characterizing the neurotranscriptomic states in alternative stress coping styles. BMC Genomics, 16(1), 425. https://doi.org/10.1186/s12864-015-1626-x

Wong, R. Y., McLeod, M. M., & Godwin, J. (2014). Limited sex-biased neural gene expression patterns across strains in Zebrafish (Danio rerio). BMC Genomics, 15(1). https://doi.org/10.1186/1471-2164-15-905

Wong, R. Y., Oxendine, S. E., & Godwin, J. (2013). Behavioral and neurogenomic transcriptome changes in wild-derived zebrafish with fluoxetine treatment. BMC Genomics, 14(1), 348. https://doi.org/10.1186/1471-2164-14-348

Wong, R. Y., Perrin, F., Oxendine, S. E., Kezios, Z. D., Sawyer, S., Zhou, L., … Godwin, J. (2012). Comparing behavioral responses across multiple assays of stress and anxiety in zebrafish (Danio rerio). Behaviour, 149(10–12), 1205–1240. https://doi.org/10.1163/1568539X-00003018

